# Multiple autism genes influence GABA neuron remodeling via distinct developmental trajectories

**DOI:** 10.1101/2025.06.25.661546

**Authors:** Kristi Zoga, Sophia Villiere, Vina Tikiyani, Andrea F. Edwards-Cintron, Pranav Thokachichu, Patrick Nicodemus, Pablo G. Camara, Michael P. Hart

**Author notes:** Corresponding author Michael P. Hart –.

## Abstract

Variation in over 100 genes are now associated with increased risk for autism and related neurodevelopmental condition, but how this variation results in distinct and overlapping behavioral changes is still not well understood. Recent efforts have focused on screening many autism genes at once for functional and phenotypic convergence, and identified subsets that are crucial for many early steps of neurodevelopment. Few studies have screened later steps of neurodevelopment, circuit function, circuit plasticity, or behaviors. We screened twenty conserved autism-associated genes for impact on experience-dependent neuron remodeling in *C. elegans*. Loss of *unc-44/ANK2*, *set-4/KMT5B, daf-18/PTEN, gap-2/SYNGAP1,* and *chd-1/CHD8* increased, while *CACNA2D3/unc-36* decreased, neurite outgrowth of the GABAergic DVB neuron in adults. Although *daf-18/PTEN, set-4/KMD5B,* and *unc-44/ANK2* had convergent phenotypes, they arise from distinct temporal trajectories with differential impact on DVB pre-synaptic morphology. Screening for the DVB regulated spicule protraction behavior identified multiple autism genes involved, but only *unc-44/ANK2* and *CACNA2D3/unc-36* were shared between screens. Application of a metric geometry computational framework (CAJAL) to the DVB morphology dataset identified 5 additional genes that impact DVB morphology, including *unc-2/CACNA1A* and *unc-10/RIMS1,* which also significantly impacted behavior. This work defines new regulators and molecular mechanisms of experience-dependent neuron remodeling and circuit plasticity, and further links these processes with conserved autism genes. It also demonstrates the utility of using intact, behavior generating circuits in *C. elegans*, to screen for novel roles for conserved autism genes.

## INTRODUCTION

Autism is defined by characteristic behavioral changes in social communication and repetitive and restricted interests, but can also include other behavioral changes ranging from anxiety, depression, seizures, to sleep disturbances^1,2^. The molecular, neuronal, and circuit mechanisms underlying the complex behavioral changes observed across the autism spectrum, and in neurodevelopmental conditions and neurodiversity more broadly, are still not well understood^3,4^. Decades of studies have uncovered a complex genetic architecture underlying the autism spectrum, now involving variation in over one hundred genes, where each gene contributes a small increase in risk for autism^5–9^. This list of high confidence or large effect autism genes provides a foothold to understand the neuronal and circuit changes that contribute to observed behavioral changes, as well as molecular insight into human neurodevelopmental processes. Systems level analyses using gene ontology, gene expression data, and protein interaction and pathway enrichments, have grouped autism-associated genes into functional categories including, 1) synaptic (or neuronal communication), 2) gene regulatory and chromatin modifying, and 3) others (*e.*g. cytoskeletal and RNA-binding)^2,5,6,9^.

Recent efforts to further refine these functional categories have screened autism-associated genes to identify functional convergence using human postmortem tissue^5,10^, cell culture^11,12^, organoids^13,14^, and animal models^15,16^. These studies have identified important functional and phenotypic convergence amongst subsets of autism-associated genes, but most have focused on the earliest neurodevelopmental processes (*e.*g. neurogenesis, neurodifferentiation, and neuronal migration). Later steps of neurodevelopment that form and refine neuronal circuits are likely also involved in autism, including axon outgrowth, synapse formation (and maturation and pruning), and experience-dependent plasticity^3,17^. Further, it is clear that autism genes function in late neurodevelopment, and after, as evidenced by increased expression of autism genes during later processes in human neurodevelopment^5^, rescue of gene-specific phenotypes after development (*SCN1A*^18,19^, *CDKL5*^20^, *FMRP*^21–24^, and *MECP2*^25^), and induction of phenotypes with genetic perturbations after development (*e.*g. *CHD8*^26^, *NRXN1*^27,28^). Screening many genes for neuronal, circuit, and behavioral phenotypes is inherently technically challenging, but a number of studies have used *C. elegans* to screen conserved autism genes at this level^29–34^.

Neuroplasticity, inhibitory interneurons, and excitatory and inhibitory (E:I) balance have all been proposed as potential convergent circuit-based mechanisms involved in autism^35–37,17^, making it critical to understand which autism-associated genes function in these processes^38^. We study the *C. elegans* GABAergic DVB neuron, which undergoes outgrowth of axon-like neurites in males in response to experience and circuit activity^39–41^. Plasticity of the DVB neuron after development directly modifies a step of male mating behavior, spicule protraction, providing a functional readout of remodeling and circuit E:I balance^39–41^. We previously found that the conserved autism-associated gene *nrx-1/NRXN1* functions in the DVB neuron to promote neurite outgrowth and inhibit spicule protraction behavior, while the conserved autism-associated gene *nlg-1/NLGN3* functions in downstream cholinergic neurons and muscles to inhibit DVB neurite outgrowth and promote spicule protraction behavior^39^. Vertebrate GABAergic neurons can undergo similar remodeling and plasticity during and after development^42–47^, best characterized in response to manipulation of animal experience^43,45,48^. Therefore, the DVB neuron is an accessible model to screen conserved autism-associated genes for convergence on experience-dependent neuron remodeling, E:I balance, and behavior.

Here, we screened twenty conserved autism-associated genes for roles in the experience-dependent neurite outgrowth of the DVB neuron in *C. elegans,* and identified phenotypes for multiple conserved autism-associated genes. *unc-44/ANK2*, *gap-2/SYNGAP1, set-4/KMT5B, daf-18/PTEN,* and *chd-1/CHD8* function to repress neurite outgrowth, while *unc-36*/*CACNA2D3* promotes neurite outgrowth. Validation and characterization of *daf-18/PTEN, set-4/KMT5B,* and *unc-44/ANK2* identified differences in the impact on DVB presynaptic structure and when each gene impacts DVB morphology. We also screened the same genes for roles in the DVB regulated spicule protraction behavior, and found limited overlap in genes that also impact DVB morphology. Application of an unbiased cell morphology analysis algorithm, CAJAL, to the DVB morphology dataset uncovered additional genes, including two with strong behavioral phenotypes. Together, this work links multiple autism-associated genes to GABAergic neuron remodeling, and identifies differences in how and when these genes act across development and into adulthood.

## RESULTS

### Multiple conserved autism-associated genes impact experience-dependent DVB neuron remodeling

We identified the *C. elegans* orthologs of twenty autism-associated genes (**Table 1**) by curating a list of high confidence genes from SFARI gene (https://gene.sfari.org/) and other recent studies^5,6^. Conserved genes were found in *C. elegans* through protein homology searches and previously published screens of autism-associated genes in *C. elegans*^29–33,49^. While in most cases this uncovered proteins/genes with very similar functions, some are less clear (e.g. the voltage-gated sodium channel *SCN2A* search returned *C. elegans cca-1*, a putative voltage-gated calcium channel, although selectivity for calcium has not been functionally confirmed^50^). We obtained at least one loss of function allele for each *C. elegans* gene to screen for impact on DVB morphology in day 3 adult males (**Table 1**, **Figure S1A**). DVB neuron morphology was analyzed using confocal microscopy of a DVB-expressed fluorescent reporter (*lim-6^int4^* promoter with green or red fluorescent protein), creation of DVB skeletons via tracing, and quantification of total neurite length and number of neurite junctions (a proxy for the number of neurite branch points).

**Table 1.** Conservation of autism-associated genes in *C. elegans*.

| SFARI Rank | Human gene | *C. elegans* ortholog | Identity (~%) | *C. elegans* alleles | CenGen DVB expression | CenGen No. of cells |
| --- | --- | --- | --- | --- | --- | --- |
| 1 | *NRXN1* * | *nrx-1* | 27 | *wy778* | yes (low) | 105 |
| 1 | *NLGN3* * | *nlg-1* | 29 | *ok259* |  | 37 |
| 1S | *SHANK3* # | *shn-1* | 23 | *tm488* |  | 32 |
| 1 | *ANK2* | *unc-44* | 31 | *e362, e1260, e1197* | yes | 128 |
| 2 | *ARHGAP5* | *syd-1* | 23 | *ju82, ok1578* | yes | 111 |
| 1S | *CACNA1A* | *unc-2* | 41 | *e55* |  | 121 |
| 1 | *CACNA2D3* | *unc-36* | 23 | *ad698, e251* | yes | 120 |
| 1S | *CHD2* | *chd-1* | 42 | *ok2798* | yes | 115 |
| 2 | *CTNND2* | *jac-1* | 25 | *ok3000* | yes | 113 |
| 1 | *DSCAM* | *C27B7.7* | 26 | *ok2978* | N/A | N/A |
| 1S | *FOXP1* | *fkh-7* | 25 | *gk793* | yes | 73 |
| 2 | *GRIA1* | *glr-2* | 37 | *ok2342* |  | 31 |
| 1 | *GRIA2* | *glr-1* | 37 | *n2461* |  | 28 |
| 1 | *GRIN2B* | *nmr-2* | 30 | *ok3324* |  | 7 |
| 1 | *KCNQ3* | *kqt-1* | 38 | *ok413* | yes | 38 |
| 1 | *KDM5C* | *rbr-2* | 30 | *ok2544* | yes | 125 |
| 1 | *KMT5B* | *set-4* | 42 | *n4600, ok1481* | yes (low) | 104 |
| 1S | *PTEN* | *daf-18* | 30 | *e1375* | yes (low) | 71 |
| 1 | *RIMS1* | *unc-10* | 24 | *e102, md1117* | yes | 118 |
| 1S | *SCN1A* | *cca-1* | 24 | *gk30* | yes | 84 |
| 1S | *SETD5* | *set-9* | 21 | *n4949* | yes (low) | 41 |
| 1S | *SLC6A1* | *snf-11* | 50 | *ok156* |  | 9 |
| 1S | *SYNGAP1* | *gap-2* | 33 | *tm748* | yes | 65 |

\* Hart and Hobert 2018<sup>39</sup>

DVB neurite length and number of neurite junctions for each gene/allele screened were normalized to the mean of controls analyzed in parallel (control mean set to 1)(**Figure 1**). The screening analysis identified 6 genes impacting DVB morphology (either one or both morphology parameters)(**Figure 1A-C**). We queried a database of hermaphrodite *C. elegans* gene expression (CENGEN)^51^ for each conserved autism-associated gene in the DVB neuron and neurons more broadly (**Table 1**). Although expression may differ between hermaphrodite and male DVB neurons, we found that many genes with no impact on DVB morphology are either lowly or not expressed in DVB, with many having broad expression across the nervous system (**Table 1**). Our screening approach identified multiple candidate autism-associated genes that regulate the experience-dependent remodeling of a GABAergic neuron.

**Figure 1.**
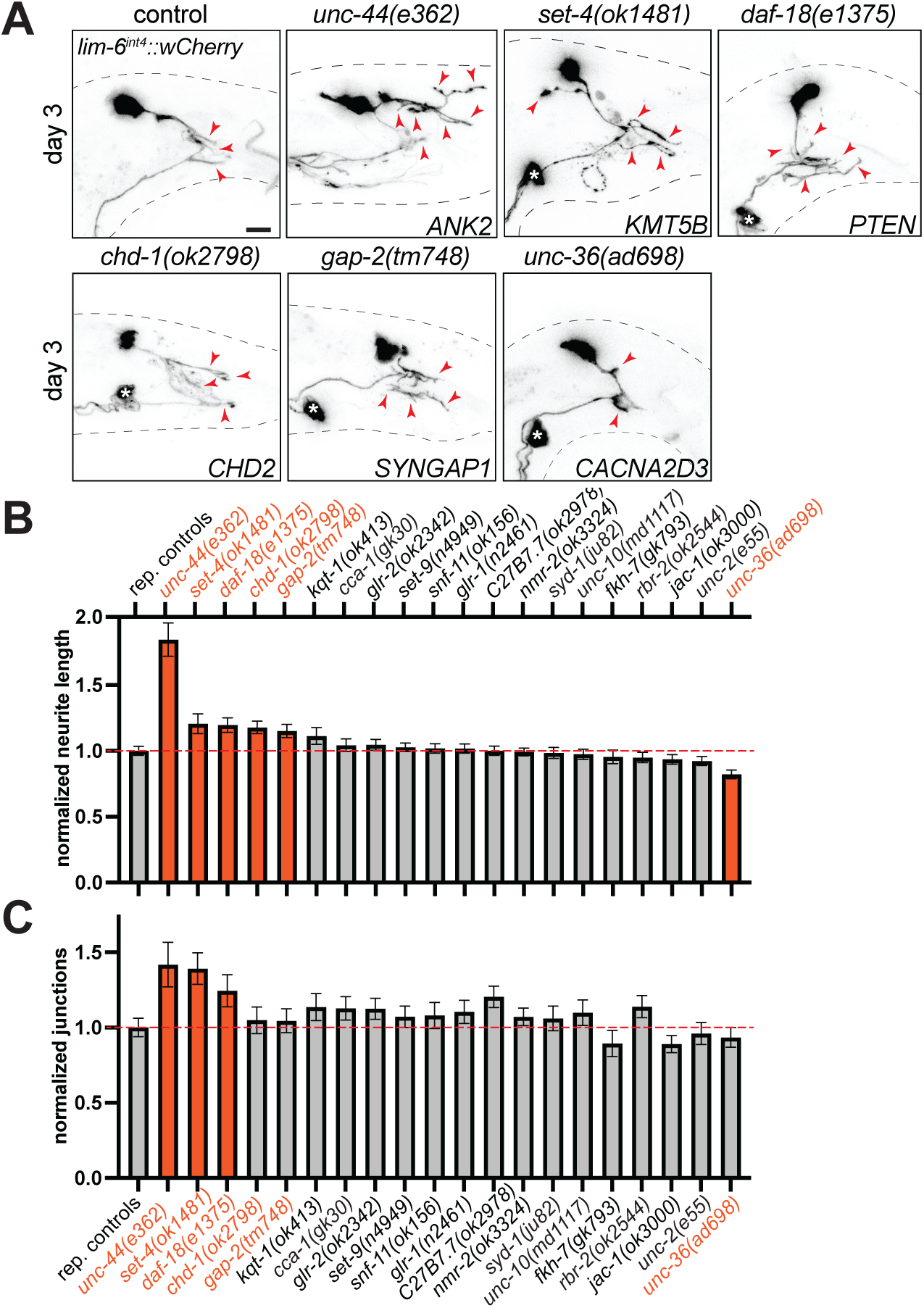
Screen of conserved autism-associated genes using DVB morphology in day 3 adult male *C. elegans*. **A)** Representative confocal micrographs of control and alleles of genes identified to impact DVB morphology in day 3 adult males (Black dashed lines represent body outline, asterisk marks PVT neuron soma (not traced, see^39^), red arrowheads indicate DVB neurite branches, scale bar = 10 μm). Quantification of normalized **B)** total neurite length, **C)** number of junctions in day 3 adult males of genotypes indicated (Bars are mean and error bars are standard error of the mean (S.E.M.), representative control group is shown for comparison, raw values for all of the normalized data and controls are included throughout the following figures or **Figure S2**). Orange bars = genes impacting DVB morphology parameters.

### DAF-18/PTEN inhibits DVB neurite outgrowth just as plasticity begins in adulthood

We next sought to validate and provide additional characterization for the top candidate genes identified in the DVB morphology screen. For this we focused on the genes that impacted both neurite length and number of neurite junctions (**Figure 1**). Mutation of *daf-18*(*e1375)*(insertion/deletion and early stop codon) increased DVB neurite length and number of junctions at day 3 of adulthood compared to controls (**Figure 2A-D**). Interestingly, when we analyzed DVB neuron morphology in day 1 adult males, *daf-*18(*e1375)* males had increased neurite length and junctions compared to control males (**Figure 2E&F**), indicating that *daf-*18 plays an earlier role in the morphology of the DVB neuron. *daf-18* may be required for development of normal morphology of DVB prior to experience-dependent plasticity in adulthood, and we therefore examined DVB neuron morphology of *daf-18* in the last larval stage before adulthood (L4) in both sexes. We found no evidence of ectopic neurite outgrowth in L4 or adult hermaphrodites (**Figure 2G&H**), indicating that development of the DVB dendrite/axon and overall morphology are not impacted by loss of *daf-18*. However, in *daf-18* L4 males we found evidence of premature DVB neurite outgrowth compared to L4 male controls (**Figure 2 G&H**). Therefore, *daf-18* inhibits DVB neurite outgrowth in L4 males, but this is likely in the context of a circuit that is beginning to undergo plasticity, representing premature DVB remodeling and neurite outgrowth. The neuronal gene expression database suggests that *daf-18* is likely expressed in hermaphrodite DVB neuron at low levels, but is expressed rather broadly in other neurons as well (**Table 1**).

**Figure 2.**
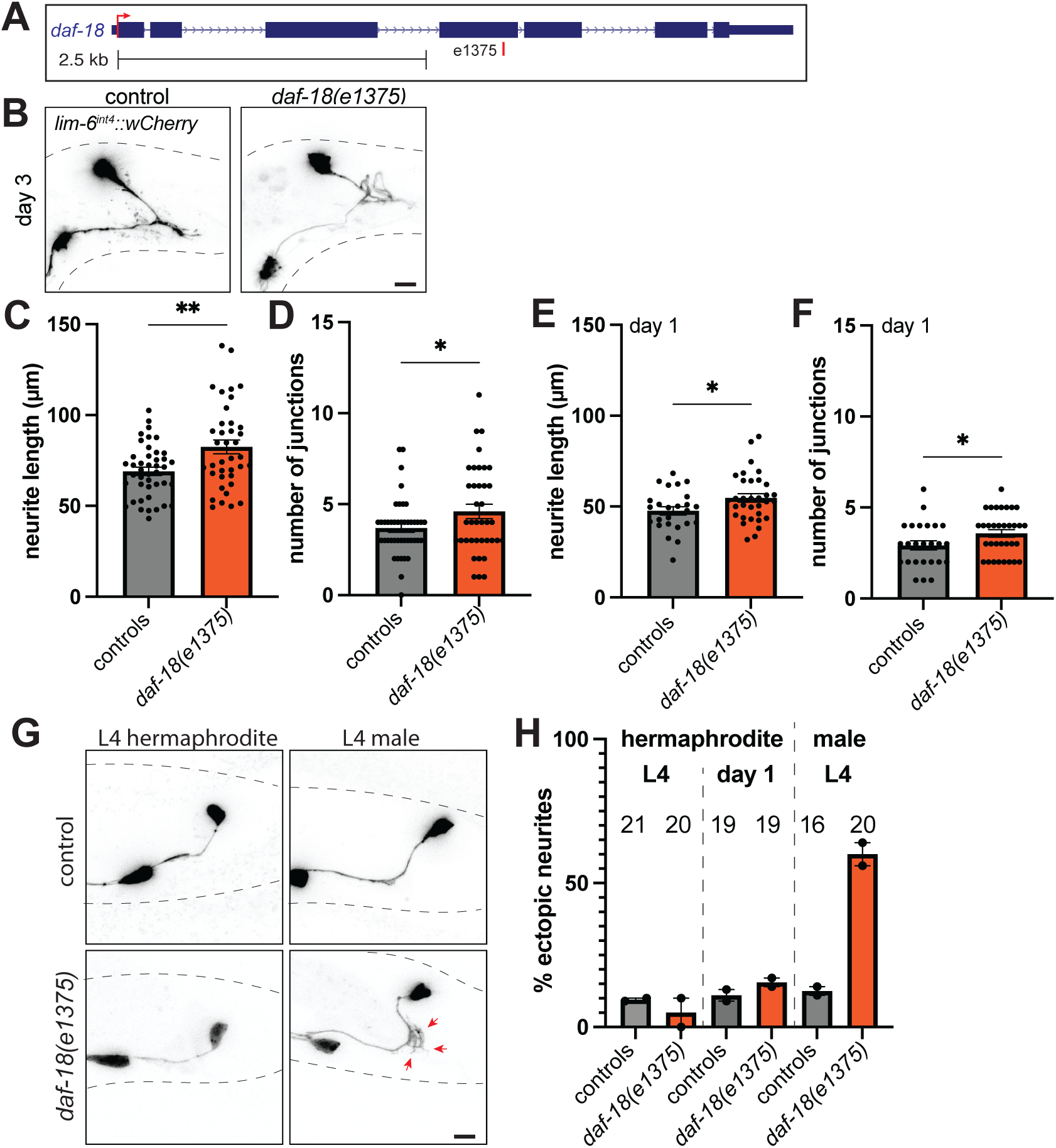
*daf-18* restricts DVB adult remodeling and developmental neurite outgrowth. A) Diagram of the *daf-18* gene locus with location of *e1375* allele shown. **B)** Representative confocal micrographs of *daf-18* and control males at day 3 of adulthood. Quantification in *daf-18* and control males at day 3 of **C)** total neurite length and **D)** number of junctions. Quantification of morphology of DVB of *daf-18* and controls in day 1 adult males of **E)** total neurite length and **F)** number of junctions. **G)** Representative confocal micrographs of DVB (*lim-6^int4^::wCherry*) in L4 hermaphrodite and L4 male (red arrowheads indicate ectopic DVB neurites). **H)** Quantification of percent of L4 and day 1 adult hermaphrodites and L4 males with ectopic neurites.

### SET-4/KMT5B inhibits DVB adulthood experience-dependent plasticity

The histone H4K20 di- and tri-methyltransferase^53^, *set-4*, had a similar impact as *daf-18* on DVB morphology at day 3 (**Figure 1**). Two loss of function alleles in *set-4, n4600*(∼1140 bp deletion) and *ok1481* (∼910 bp deletion), increased DVB neurite length and number of junctions compared to controls at day 3 of adulthood (**Figure 3A-D**). *set-4* is expressed in the nervous system and other tissues in *C. elegans*, with evidence of low expression in the hermaphrodite DVB neuron (**Table 1**). We also analyzed the impact of *set-4* alleles at day 1 and found no changes in DVB morphology compared to controls, while at day 5 we found a significant increase in DVB neurite length in *set-4* compared to controls (**Figure S3A-E**). This supports a role for *set-4* in DVB morphology and remodeling specifically in adulthood. Lastly, we analyzed DVB pre-synaptic puncta in *set-4* alleles and found no change in the number of *cla-1* pre-synaptic puncta or area at day 3 of adulthood compared to controls (**Figure 3E-G**). However, in the context of the increased DVB neurite length observed in *set-4* (**Figure 3B&C**), this indicates a potential synaptic connectivity phenotype with altered distribution of active zones along DVB neurites.

**Figure 3.**
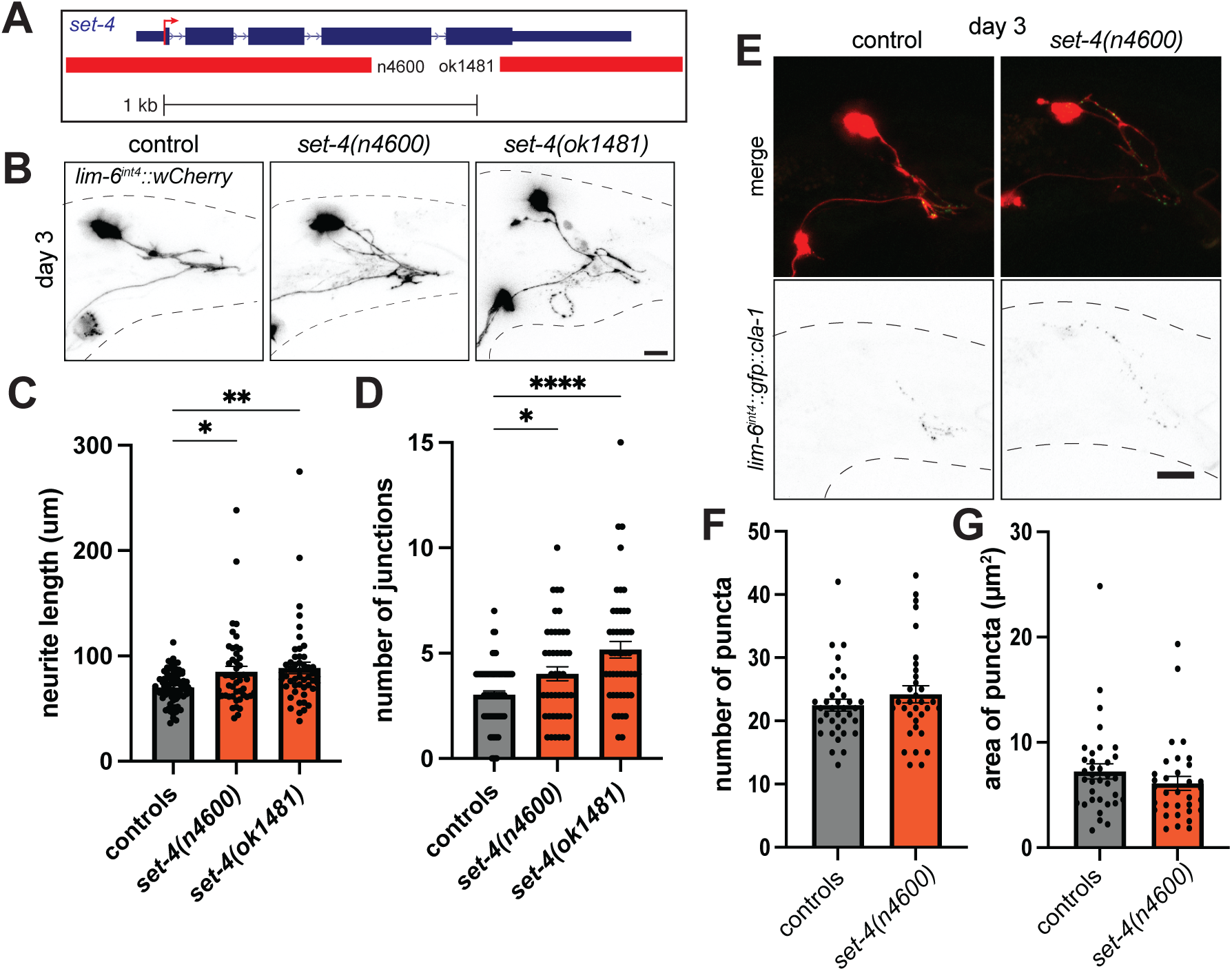
*set-4* restricts DVB neurite outgrowth in adult males. A) Diagram of the *set-4* gene locus with location of included alleles shown. **B)** Representative confocal micrographs of *set-4* alleles indicated and controls in day 3 adult males. Quantification of morphology of DVB of *set-4* and controls in day 3 adult males of **C)** total neurite length and **D)** number of junctions. **E)** Representative confocal micrographs of DVB expressed GFP-tagged CLA-1 (*lim-6^int4^::gfp::cla-1*) in day 3 adult *set-4* and control males. Quantification of number of **F)** synaptic puncta and **G)** area of puncta.

### UNC-44/ANK2 inhibits DVB neurite outgrowth in development and adulthood

Our screen also identified that the ankyrin repeat containing gene, *unc-44,* plays a role in DVB neurite length and junctions (**Figure 1**). Alleles impacting the long isoform of *unc-44* (*e362* 1bp sub early stop, and *e1260*, not curated but predicted to be in long isoform exon) significantly increased DVB neurite length and number of neurite junctions at day 3 (**Figure 4A-D**). We observed a similar impact on DVB morphology in a third allele, *e1197*, but did not quantify DVB morphology as the frailty of these animals made it difficult to obtain a satisfactory number of animals for microscopy (**Figure 4B**). We next analyzed the impact of *unc-44(e362)* on DVB at day 1 of adulthood and found a significant increase in DVB neurite outgrowth compared to controls (**Figure 4E-F**). While we find a role for *unc-44* in adult remodeling of DVB, *unc-44* has known roles in axon guidance and branching in neurons during development, including roles in the complex PVD sensory neuron dendritic arbor^54,33,55^.

**Figure 4.**
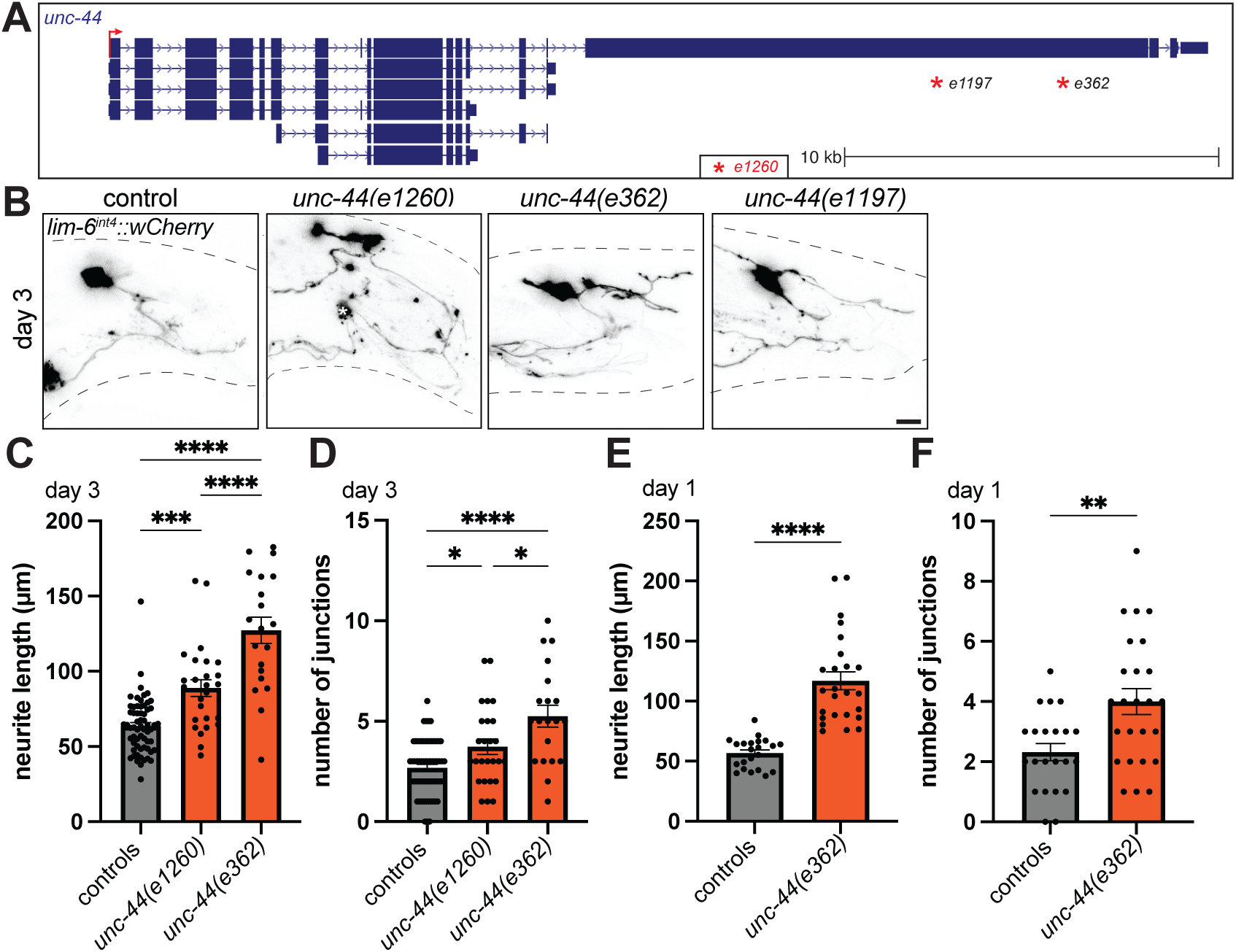
*unc-44* restricts DVB neurite outgrowth in adulthood. A) Diagram of the *unc-44* gene locus with location of alleles shown (e1260, premature truncation, but location not curated^33^). **B)** Representative confocal micrographs of *unc-44* alleles indicated and controls in day 3 adult males. Quantification of morphology of DVB of *unc-44* and control day 3 adult males of **C)** total neurite length and **D)** number of junctions. Quantification of morphology of DVB of *unc-44* and controls in day 1 adult males of **E)** total neurite length and **F)** number of junctions.

We therefore examined DVB morphology during the last larval stage (L4) and in adult hermaphrodites and found ectopic DVB neurites at L4 in both sexes and in adult hermaphrodites (**Figure 5A-B**). We also examined L2 animals, and found ectopic DVB neurites in *unc-44* at this earlier larval stage (**Figure 5A-B**). The small ectopic neurites observed in developing and adult hermaphrodite animals are not at the level of neurite outgrowth observed in adult control of *unc-44* mutant males. These results implicate the long isoform of *unc-44* in the development of DVB axon morphology, which may be similar or distinct to its role in experience-dependent remodeling in adulthood. These strong phenotypes led us to also analyze DVB pre-synaptic morphology with loss of *unc-44*. We found that *unc-44(e362)* mutant males had increased number and area of DVB presynaptic puncta at day 3 of adulthood compared to controls (**Figure 5C-E**). Therefore, *unc-44* functions during development of DVB to inhibit a limited number of axon branches, and appears to play an even larger role in stabilizing and inhibiting DVB experience-dependent neurite outgrowth after development.

**Figure 5.**
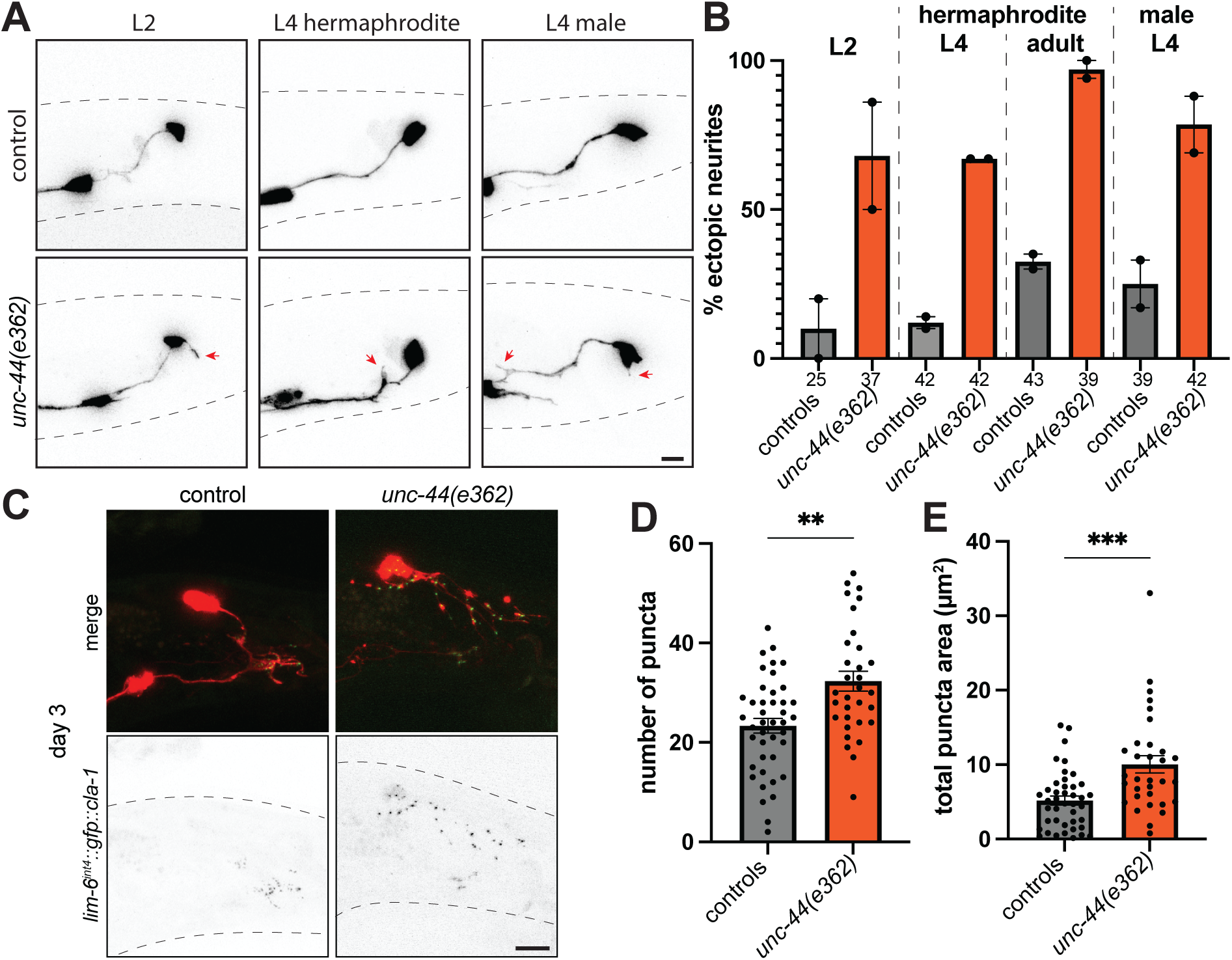
*unc-44* is required for development of DVB morphology and pre-synaptic refinement. A) Representative confocal micrographs of DVB (*lim-6^int4^::wCherry*) in L2 and L4 hermaphrodite, and L4 male (ectopic neurites indicated by red arrowheads). **B)** Quantification of percent of L2, L4, and day 1 adult hermaphrodites and L4 males with ectopic DVB neurites. **C)** Representative confocal micrographs of DVB expressed GFP-tagged CLA-1 (*lim-6^int4^::gfp::cla-1*)in day 3 adult *unc-44* and control males. Quantification of number of **D)** synaptic puncta and **E)** area of puncta.

### Conserved autism-associated genes impact spicule protraction behavior

We also screened the same conserved autism-associated genes for impact on the behavior impacted by DVB plasticity, spicule protraction, which is a crucial step of male mating required for sperm transfer. To analyze spicule protraction behavior, we quantified the time for males to fully protract their spicules when exposed to the acetylcholinesterase inhibitor aldicarb (**Figure S1**). This assay is an established model for inhibitory and excitatory balance at neuromuscular junctions^56,57^ and in the spicule protraction circuit^39^. We normalized time to spicule protraction for each gene to their respective controls analyzed in parallel, with the mean of the controls set to 1 (**Figure 6A**). Interestingly, most of the genes impacting spicule protraction behavior were not the same as those impacting DVB neurite length and junctions (**Figure 1&6A&7**), and a fisher’s exact test indicates no significant relation (odds ratio = 0.90, p-value = 1) between genes impacting each phenotype. To this point, we found no significant impact on spicule protraction behavior in day 3 *daf-18* and *set-4* males, or day 1 *daf-18* males (**Figure S3F-H**).

**Figure 6.**
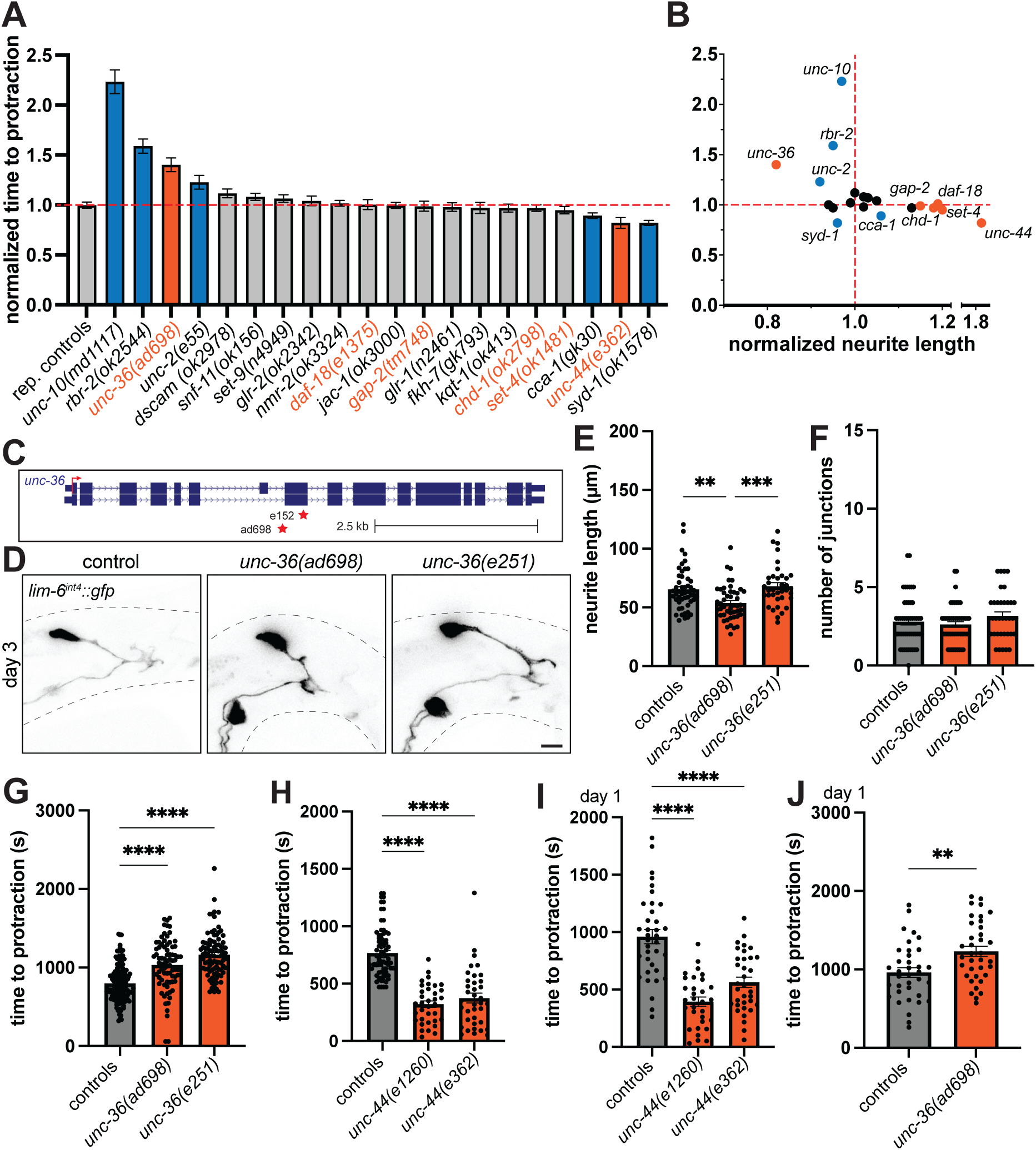
Loss of conserved autism-associated genes alters spicule protraction behavior. **A)** Normalized time to spicule protraction behavior on aldicarb in day 3 adult males of genotypes indicated (Bars are mean and error bars are standard error of the mean (S.E.M.), representative control group is shown for comparison, raw values for all the normalized data and controls are included in other figures). Orange bars = genes impacting DVB morphology and spicule protraction behavior, blue bars = genes impacting spicule protraction behavior. **B)** XY plot of normalized time to spicule protraction and normalized neurite length (from Figure 1) for each gene. **C)** Diagram of the *unc-36* gene locus with location of alleles annotated. **D)** Representative confocal micrographs of DVB in day 3 control and *unc-36* males (Black dashed lines represent approximate body outline, scale bar = 10 μm). Quantification of **E)** DVB total neurite length and **F)** number of junctions, and **G)** time to protraction in day 3 control and *unc-36* males. **H)** Quantification of time to protraction in day 3 control and *unc-44* males. Quantification of time to spicule protraction of day 1 **I)** *unc-44*, **J)** *unc-36*, and respective control males.

Despite lack of overlap between genes impacting DVB morphology and behavior, we did find genes that impacted both phenotypes. Mutation in the voltage-gated calcium channel subunit *unc-36*(*ad698)* resulted in a small, but significant, decrease in DVB neurite length in day 3 adult males compared to controls (there was no significant change in DVB morphology observed in *unc-36*(*e152*))(**Figure 6C-F**). *unc-36* loss resulted in a significant increase in the time to spicule protraction at day 3 compared to controls (*ad698* and *e152,* both 1bp substitutions and early stop alleles^58^)(**Figure 6G**). *unc-44* also impacted both DVB morphology and spicule protraction behavior (**Figure 6G**). Loss of *unc-44* (*e362* 1bp sub early stop, and *e1260*, not curated) significantly decreased the time to spicule protraction at day 3 compared to controls (**Figure 6H**). With the large impact of *unc-36* and *unc-44* on spicule protraction behavior at day 3, we analyzed day 1 adult males and found that both genes also impacted the time to spicule protraction compared to controls (**Figure 6I-J**). For *unc-44*, where we found an increase in the number of presynaptic active zones (**Figure 5C-E**), it seems likely that these additional synapses are likely dysfunctional or connect with targets outside the protraction circuit based on the opposite behavioral phenotype than expected. Perhaps a reduction in GABAergic inhibition in the circuit occurs as additional ectopic connections lower functional connectivity to inhibit spicule protraction targets.

We found two genes that had large impacts on spicule protraction behavior in opposite directions (**Figure 6A**), but with no significant impact on neurite length or junctions. Mutation in the small GTPase *unc-10* (*md1117*, ∼8000 bp deletion and/or *e102*, 1bp substitution) did not impact DVB neurite length of junctions at day 3 (**Figure 7A,C-E**), but drastically increased the time to spicule protraction compared to controls at day 1 and 3 (**Figure 7H-I**). Mutation of the Rho GTPase homolog *syd-1*(*ok1578)*(∼700 bp deletion) also did not impact DVB neurite length of junctions at day 3 (**Figure 7B,C,F,&G**), but decreased the time to spicule protraction on aldicarb compared to control males at day 3 of adulthood (**Figure 7J**). A milder allele of *syd-1(ju42)*(1 bp substitution) had no impact on morphology or the time to spicule protraction compared to controls (**Figure 7B&J**). Both *unc-10* and *syd-1* are expressed broadly in many neurons and may be expressed in DVB (**Table 1**)^51^. These two GTPases have opposite impacts on spicule protraction, with *unc-10* potentially reducing cholinergic output and *syd-1* potentially reducing GABA output from the DVB neuron.

**Figure 7.**
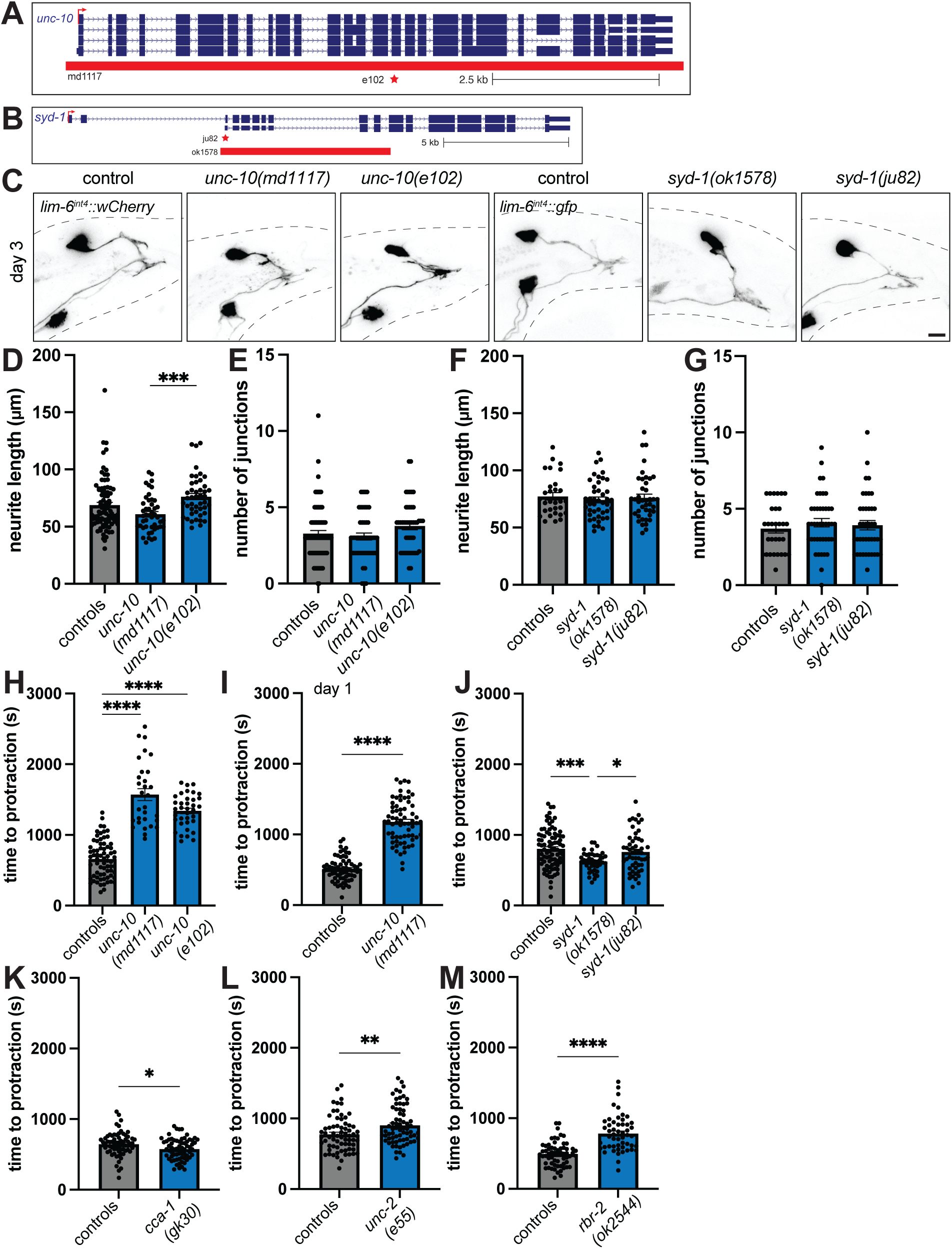
Spicule protraction behavior is impacted by *unc-10, syd-1, cca-1, unc-2*, and *rbr-2.* **A)** Diagram of the *unc-10* gene locus with location of alleles shown. **B)** Diagram of the *syd-1* gene locus with location of alleles shown. **C)** Representative confocal micrographs of *unc-10 and syd-1* alleles indicated and controls in day 3 adult males. Quantification of morphology of DVB of *unc-10 and syd-1* and controls in day 3 adult males of **D/F)** total neurite length, **E/G)** number of junctions. Quantification of time to spicule protraction on aldicarb of *unc-10* and respective control males at **H)** day 3 or **I)** day 1 of adulthood. Quantification of time to spicule protraction on aldicarb at day 3 of adulthood of control and **J)** *syd-1*, **K)** *cca-1*, **L)** *unc-2*, **M)** *rbr-2* males.

Mutation of the low voltage-gated calcium channel alpha subunit *cca-1*(*gk30)*(∼260 bp deletion) decreased the time to spicule protraction on aldicarb compared to control males at day 3 of adulthood (**Figure 7K**). Mutation of the high voltage-gated calcium channel *unc-2*(*e55)*(substitution introducing an early stop codon), and the histone demethylase *rbr-2*(*ok2544)*(∼1300 bp deletion), increased the time to spicule protraction on aldicarb compared to controls at day 3 (**Figure 7L&M**). *unc-2* is likely not expressed in DVB, while *cca-1* and *rbr-2* may be expressed in DVB, although *rbr-2* is expressed in broadly in many neurons (**Table 1**)^51^. Overall we find that conserved autism genes involved in calcium signaling and calcium regulation at the synapse are crucial for spicule protraction behavior induced by aldicarb, and interestingly *rbr-2* also impacts spicule protraction behavior but has very different function.

### CAJAL confirms and expands conserved autism-associated genes with roles in DVB morphology

We recently reported the development and use of CAJAL to analyze neuron morphologies, including in the DVB neuron in *C. elegans;* where CAJAL confirmed previously identified conserved autism-associated gene phenotypes (*nrx-1*, *nlg-1*, and *shn-1*)(**Table 1**)^52^. CAJAL is a computational framework that uses metric geometry to compare cell morphologies, where distances between points in the morphology latent space indicate the physical deformation necessary to change the morphology of one neuron into another. CAJAL compares neuron morphologies without bias towards any specific morphological parameters (e.g., neurite length or number of branches), analyzing the overall shape of the neurons. The DVB neuron is somewhat unique in *C. elegans* in that it undergoes structural plasticity and its morphology is variable between animals^39^. We used CAJAL first to compare traced skeletons of the DVB neuron from day 1 and day 3 adult males to traced skeletons of a less plastic neuron in hermaphrodite *C. elegans*, the RIC octopamine-producing interneurons (Left and Right). First, we find that the diversity of neuron morphologies is substantially larger in DVB neurons on day 3 than in RIC neurons at day 3, as expected (**Figure 8A-B**). The simple morphology of day 1 DVB shows decreased morphologic diversity compared to RIC, but day 3 DVB exhibits increasingly diverse morphologies from day 1 to day 3 (**Figure 8A-B**), further confirming the morphologic plasticity of DVB and potential for CAJAL to identify differences in morphology of the DVB and other *C. elegans* neurons.

**Figure 8.**
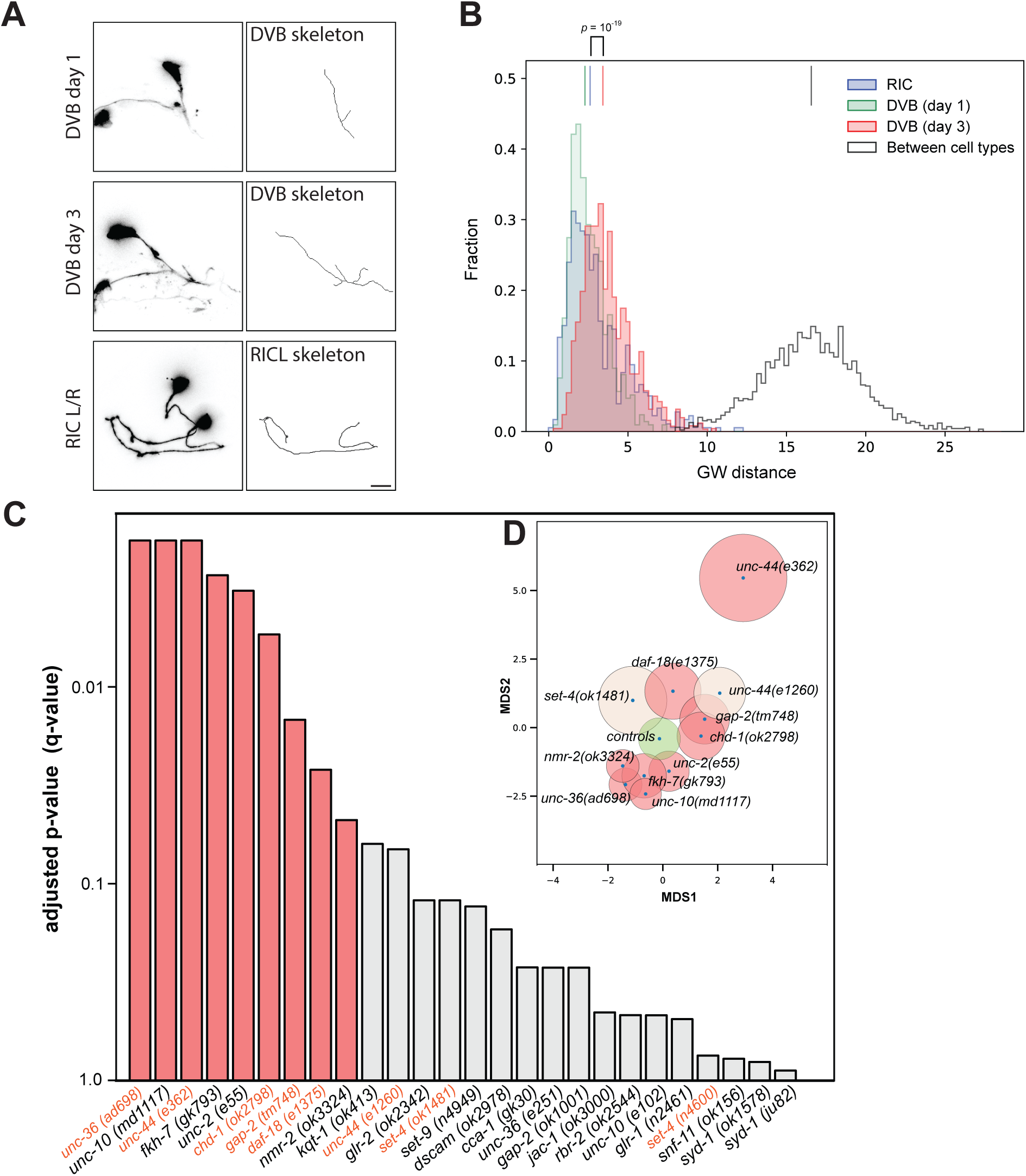
CAJAL algorithm confirms and identifies additional autism-associated genes involved in DVB morphology. **A)** Representative confocal micrographs and skeleton traces of the DVB neuron in day 1 and 3 adult control males and the RIC neuron in day 3 adult control hermaphrodites. **B)** Histogram of GW pairwise distances for DVB and RIC neuron trace skeletons (vertical lines indicate the medians). The difference between RIC and DVB on day 1 is also significant but much smaller (two-tailed Wilcoxon rank-sum p-value = 10^-4^)). **C)** Barplot showing the significance of each mutation from adjusted p-value (q-value). **D)** A low-dimensional MDS skeleton of the morphology space for the controls and significant mutations, where the diameter of each circle indicates the amount of morphological variability within each population (and corresponds to ½ of the standard deviation in the pairwise GW distance) and the distance between blue points indicate the GW distance between centroids.

Therefore, we applied CAJAL to compare the traced skeletons and morphology of the DVB neuron between controls and the screened conserved autism-associated genes, hoping to identify changes in morphology beyond the simple parameters of neurite length and number of junctions. CAJAL identified 9 genes/alleles where DVB morphology was significantly different from the DVB morphology of the combined controls (**Figure 8C**). This included 5 of the 6 genes identified analyzing just 2 morphologic parameters, with *set-4(ok1481)* having an adjusted p-value (q-value) of 0.13 and, interestingly, a morphological variability that is substantially larger than that of controls and other gene alleles. CAJAL identified 4 additional genes not identified by the 2 screening parameters, including *unc-2* and *unc-10,* which both had significant impacts on spicule protraction behavior (**Figure 8C**, **Figure 6A&B**). Two genes identified by CAJAL that did not significantly impact morphology or behavior in our screening were the NMDA-like glutamate receptor, *nmr-2,* and forkhead transcription factor, *fkh-7* (**Figure 8C**). We also plotted morphological representatives (centroids) of the controls and the significant gene/alleles (plus *unc-44(e1260)* (adjusted p-value = 0.06) and *set-4(ok1481)*) in a low-dimensional multidimensional scaling (MDS) representation of the morphology space to visualize the amount of morphological variability (diameter of circles) and impact on morphology (space between centroids – dots) within each population (**Figure 8D**). This plot demonstrates that genes/alleles can impact DVB morphology in different ways. For example, the morphology of DVB in *unc-44* alleles is very different than in *unc-10* alleles, but DVB morphology is not very different between *unc-10* and *unc-36* alleles (**Figure 8D, Figure S4**), which can be visualized by comparing centroid neurons of each population (**Figure S4**). Some genes, like *unc-44* and *set-4,* also lead to a larger morphologic variability than observed in the controls (**Figure 8D, Figure S4**). It is possible that genes/alleles with overlap and closer centroids may function in similar ways or pathways to impact DVB morphology in similar ways.

### Directional impact of autism-associated on neurite outgrowth predicts distinct synaptic compartment localization

We next asked whether the genes identified and grouped in the DVB *C. elegans* screen might provide additional context for their synaptic or neuronal functions. We compiled a list of human orthologs of the *C. elegans* genes included in our screening study with the addition of previously identified genes (*nrx-1/NRXN1, nlg-1/NLGN3,* and *shn-1/SHANK3*)^39,52^(**Supplemental Table 1**). The list of human autism-associated genes includes 50 genes as a number of *C. elegans* genes are the singular ortholog of a family of human genes (e.g. *nrx-1* to *NRXN1-3*) or have similar identity to more than one human autism-associated gene. We then used SynGO (<u>Syn</u>aptic <u>G</u>ene <u>O</u>ntologies)^59^ on the human genes grouped by phenotypes for DVB morphology and spicule protraction behavior (from **Supplemental Table 1**). SynGO identified enrichments for synaptic genes generally, as expected, with a mix of pre- and post-synaptic location and functional ontology terms (**Figure 9A&B, Supplemental Table 2**). We then separated genes by the approximate directional impact on DVB neurite outgrowth. The genes that increased DVB neurite outgrowth showed enrichment of ontology terms for postsynaptic specialization, integral components of post synapse, and postsynaptic density, while the genes that decreased DVB neurites were enriched for ontology terms for presynaptic active zone, integral components of presynaptic active zone and membrane, and pre-synapse (**Figure 9A, Supplemental Table 2**). Therefore, there are preliminary differences in the synaptic functional and location terms enriched for the genes grouped by their DVB phenotypes in *C. elegans* and may provide insight into convergence of autism-associated genes on neuron remodeling in other systems.

**Figure 9.**
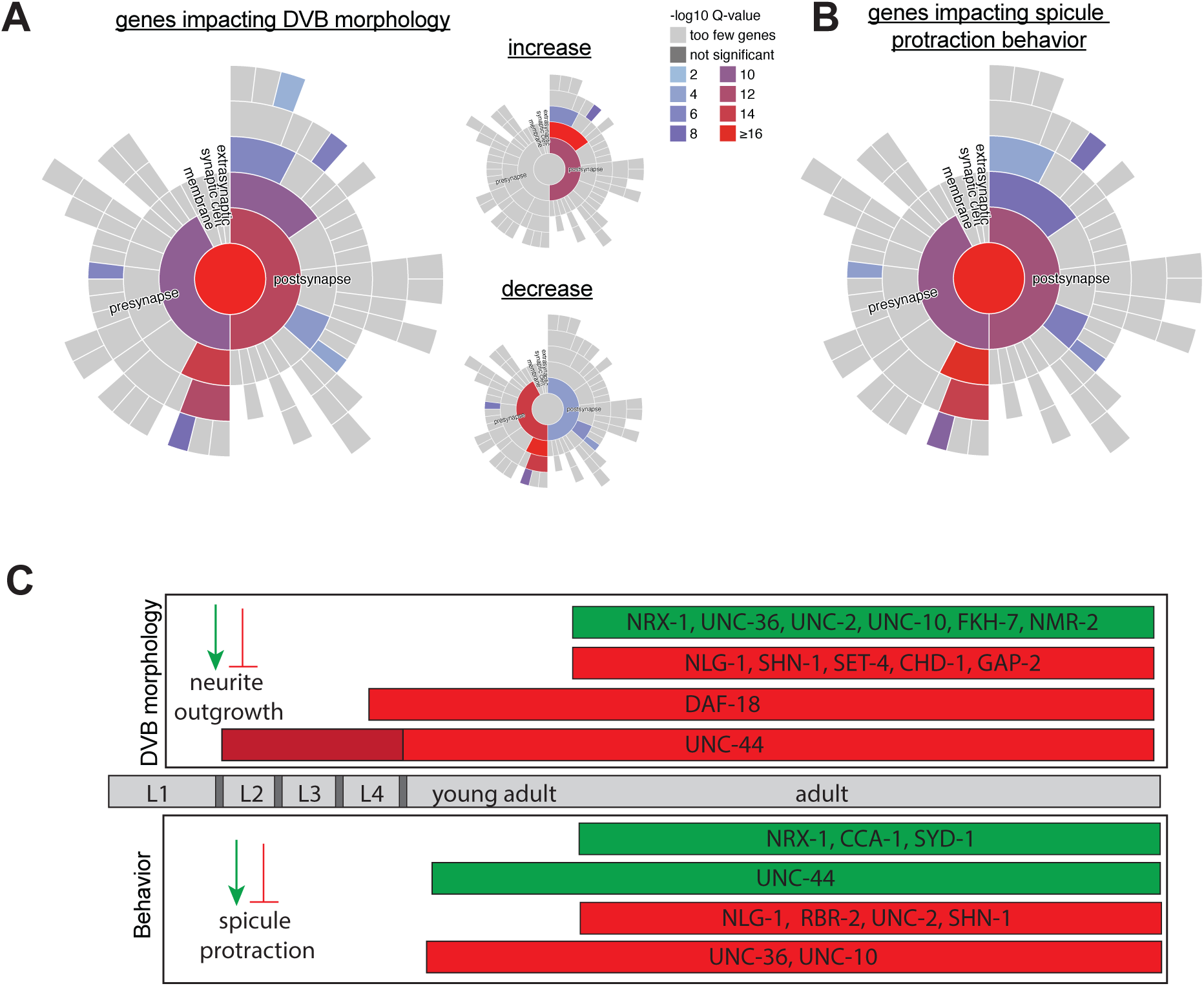
Synaptic Gene ontology and temporal model for autism-associated genes in DVB remodeling. <u>Syn</u>aptic <u>G</u>ene <u>O</u>ntology analysis of genes identified in our screen and by CAJAL was performed with the use of SynGO^59^. Sunburst plots for synaptic location term enrichments for genes impacting **A)** DVB morphology (insets show sunburst for genes that generally either increase or decrease DVB neurite outgrowth), or **B)** spicule protraction behavior (sunburst plots showing first level terms of pre-synapse and post-synapse, second level terms include specialization, active zone, and membrane). **C)** Cartoon timeline of *C. elegans* developmental stages into adulthood (gray). Genes are placed in bars indicating timing and directionality of function on DVB neurite outgrowth or aldicarb-induced spicule protraction behavior based on the phenotypes observed in loss of function alleles for each gene (green = promote, red = inhibit).

## DISCUSSION

Using DVB neuron remodeling in *C. elegans*, we identified roles for multiple conserved autism-associated genes in regulating GABAergic neuron remodeling, circuit connectivity, E:I balance, and behavior. The majority of the genes identified in our screen function to restrict experience-dependent neurite outgrowth, including both synaptic (*unc-44/ANK2*, *gap-2/SYNGAP1*, and *daf-18/PTEN*) and chromatin modifying genes (*set-4/KMT5B* and *chd-1/CHD8*)(**Figure 9C**). Analysis of the DVB dataset beyond simple morphologic parameters, using CAJAL, confirmed and expanded the list of genes that impact DVB morphology. Altogether, these analyses identified 13 genes that fall into three phenotypic groups, and importantly 7 genes with no significant phenotypes. *unc-36/CACNA2D3, unc-44/ANK2, unc-10/RIMS,* and *unc-2/CACNA1A* impacted DVB morphology and spicule protraction behavior; *gap-2/SYNGAP1*, *daf-18/PTEN, set-4/KMT5B,* and *chd-1/CHD8* impacted only DVB morphology (CAJAL also identified *fkh-7/FOXP1* and *nmr-2/GRIN2B*); and *cca-1/CACNA1H, syd-1/ARHGAP5,* and *rbr-2/KDM5B* impacted only spicule protraction behavior (**Supplementary Table 1**). Synaptic gene ontology of conserved genes grouped by directional impact on DVB neurite outgrowth identified potential differences in pre- vs. post-synaptic localization in humans. Our results identify convergence of conserved autism-associated genes in GABAergic neuron remodeling, building a framework to study how, when, and where genes function and interact in the DVB circuit and other models of experience-dependent neuron remodeling.

We validated and characterized the genes that most significantly increased DVB neurite length and junctions, including *daf-18/PTEN, set-4/KMT5B,* and *unc-44/ANK2.* All three genes similarly impacted DVB experience-dependent remodeling at day 3 of adulthood, but *daf-18* resulted in ectopic neurites in L4 males and *unc-44* resulted in ectopic neurites even earlier in development in both sexes. Therefore, the convergent DVB morphology phenotype observed is due to distinct temporal impact and mechanisms for each gene. This is even more striking in the context of our previous work on *nrx-1/NRXN1* and *nlg-1/NLGN3*, which impacted DVB morphology at and after day 3^39,52^, similar to *set-4/KMT5B.* Only *unc-44/ANK2* resulted in altered pre-synaptic morphology and a behavioral change, although *set-4/KMT5B* also displayed a subtle synaptic number/area change in the context of increased neurite length. The discrepancy between morphology and behavioral changes observed for *set-4/KMT5B* (and other genes) could be specific to the aldicarb assay. Although this assay was previously correlated with mating induced spicule protraction and mating efficiency^39^, it is a pharmacologic manipulation that differs significantly from spicule protraction during a natural mating interaction. Genes without a behavioral phenotype could also impact the DVB neuron and circuit in ways that are only evident with a second hit, for example upon stress or sleep deprivation, which we found alter DVB remodeling and behavior in a genotype-specific manner^40,41^. Overall, these results demonstrate the utility of the DVB model, which allows comparisons between genes, timepoints, and sexes to identify specific functions for genes in neurite outgrowth occurring in development, adulthood, or both.

The phenotypes observed in our morphology and behavior screens have similarities and discrepancies with reported gene functions across model systems and assays. Consistent with our results, *unc-44/ANK2* is important for development and maintenance of neuron morphology across multiple neuron types and animal models^33,54,60–64^, while *daf-18/PTEN* impacts axons and dendrites with notable differences between neuron subtypes and organisms^65,67^. Knockdown of *set-4/KMT5B* decreased dendritic complexity and increased dendritic spine density in cultured mouse neurons^68^, and *fkh-7/FOXP1* and *chd-1/CHD8/2* impacted axon and dendrite morphogenesis and E:I balance^69–71^. However, *syd-1/ARHGAP5* impacts developmental neurite outgrowth of GABAergic motor neurons in *C. elegans*, suggesting that DVB outgrowth in adult males (not impacted by *syd-1*) is mechanistically distinct. Further, while we observed *syd-1* to impact aldicarb induced spicule protraction, it did not impact aldicarb induced muscle paralysis^57^, indicating circuit specificity or a technical differences between studies (RNAi versus loss of function allele). The spicule protraction phenotypes of calcium signaling genes (*unc-2*, *unc-36*, and *unc-10*) are consistent with roles in aldicarb induced body wall muscle paralysis^57^, suggesting conserved involvement across *C. elegans* neuromuscular junctions and behaviors. Lastly, *unc-36/CACNA2D3, unc-2/CACNA1A, syd-1/ARHGAP5, unc-10/RIMS1,* and *nrx-1/NRXN1* all interact and alter synapse function and size in *C. elegans* and mice in a context specific manner^72–76^. Therefore, the functions of many genes suggest conservation across neuron types, behaviors, and even organisms, but clearly the temporal and neuronal context is crucial when comparing gene phenotypes.

With the recent focus on screening for functional and phenotypic convergence amongst autism genes, comparing results is crucial to understand the utility of this approach. We compared the screening results reported here (including *nrx-1*, *nlg-1*, and *shn-1*) with six other published *C. elegans* screens of autism genes, which examined different animal, neuron morphology, and behavioral phenotypes^29–34^. For genes included in at least three studies, every study observed at least one phenotype for *nrx-1* (5 screens), *nlg-1* (5 screens), *egl-19* (3 screens), *unc-2* (3 screens), and *unc-44* (3 screens), while other commonly tested genes also had high hit rates, including *daf-18* (4/5 screens), *nmr-2* (4/5 screens), and *gap-2* (3/5 screens). We uniquely included and found DVB or spicule protraction phenotypes for *chd-1* and *syd-1*, but not for *snf-11* and *glr-2*, which had phenotypes in multiple other screens (3 and 2, respectively). A clear caveat in this comparison is that genes lacking phenotypes could be false negatives due to screening parameters or due to different methods used for gene perturbation or knockdown. Together, the *C. elegans* screening studies successfully identified conserved autism-associated genes impacting multiple aspects of *C. elegans* neurobiology and behavior, including genes with convergent and restricted functions. Comprehensive comparison of functional screens for autism genes across organisms and phenotypes holds the potential to group autism genes by functions across neurodevelopmental processes, neuron and circuit types, and behaviors, as well as identifying genes with broad or restricted functions.

Many autism-associated genes are clearly linked with synaptic properties and E:I balance, suggesting together with our work, that DVB neurite outgrowth and behavior may be readout of these functions more broadly. Although DVB neuron remodeling in *C. elegans* can provide unique insights into our understanding of circuits, behaviors, and gene function, there are clear limitations connecting this work to human genes, behaviors, and autism and related conditions. Our work relies on homozygous loss of function alleles, only rarely found in autism, which includes almost every other form of genetic variation and inheritance possible^2,6,7^. Therefore, testing different types of alleles on various phenotypes is crucial to understand how each gene can contribute to any phenotype. Our results highlight the utility of using a small model with intact, behavior generating, circuits to identify functional convergence of many genes, and to uncover gene interactions and molecular mechanisms involved in complex neurobiological processes. To understand how genes converge onto behavioral and neuronal phenotypes, as is the case across the autism spectrum, will require analyzing and comparing across biological and neurodevelopmental process, cell/neuron type, and developmental stage. Even in a small nervous system with reduced complexity, we find a remarkable complexity and interplay of the functions of conserved autism-associated genes. This further highlights why despite having a plethora of genetic associations for many years, finding actionable mechanisms and interactions involved in autism and related conditions has remained elusive^2^. This work provides a list of conserved autism-associated genes with novel convergent functions and phenotypes in a complex and dynamic process that occurs late in, and after, neurodevelopment. These results provide a framework to guide future experiments on conserved genes and mechanisms underlying neuron remodeling in other organisms, which remain limited due to the technical challenges associated with studying this process^42,77,44,47,45^.

## Supplemental Figure Legends

**Supplemental Figure 1. Screening method cartoon and genomic locus diagrams. A)** Cartoons of L4 and adult male *C. elegans*, and DVB neuron morphology from L4 into early adulthood with day 3 indicated as time at which screening for DVB neuron morphology and spicule protraction behavior were performed. **B)** Cartoon of DVB and spicule protraction neurons and muscles in tail of male *C. elegans.* Aldicarb inhibits acetylcholinesterase at neuromuscular junctions of SPC and DVB onto spicule protractor muscle, allowing acetylcholine from SPC to buildup and cause spicule protractor muscle contraction in presence of aldicarb. **C)** Genomic loci diagrams for candidate genes and alleles used in the screen and in validation experiments not shown in main figures, with allele locations indicated by red bars indicating deletion and red asterisks indicating point mutations (early stop of missense mutations).

**Supplemental Figure 2. Conserved autism-associated genes screened without impact on DVB morphology or spicule protraction behavior. A)** Representative confocal micrographs of control and loss of function alleles of genes indicated in day 3 adult males with DVB reporter (*lim-6^int4^::wCherry*). Quantification of **B)** total neurite length, **C)** number of junctions, and **D)** time to spicule protraction behavior on aldicarb in day 3 adult males of genotypes indicated. **E)** Representative confocal micrographs of control and alleles of genes indicated in day 3 adult males with DVB reporter (*lim-6^int4^::gfp*). Quantification of **F)** total neurite length, **G)** number of junctions, and **H)** time to spicule protraction behavior on aldicarb in day 3 adult males of genotypes indicated.

**Supplemental Figure 3. set-4 impact on DVB morphology at day 1 and day 5. Set-4 and daf-18 do not impact spicule protraction behavior.** A) Representative confocal micrographs of *set-4* alleles indicated and controls in day 1 and day 5 adult males. Quantification of DVB in *set-4* and controls in day 1 adult males of **B)** total neurite length and **C)** number of junctions and day 5 adult males of **D)** total neurite length and **E)** number of junctions. Time to spicule protraction of controls and *daf-18* males at **F)** day 3 and **G)** day 1 of adulthood. **H)** Time to spicule protractions of control and *set-4* males at day 3 of adulthood. adult male controls.

**Supplemental Figure 4. DVB centroid examples identified by *CAJAL* for a subset of conserved autism-associated genes.** Confocal micrographs and skeleton traces of the DVB neuron in 3 adult males of the CAJAL identified centroid neuron of each population indicated in the low-dimensional morphology space (the diameter of each circle indicates the amount of morphological variability within each population (and corresponds to ½ of the standard deviation in the pairwise GW distance) and the distance between blue points indicate the GW distance between centroids) (red circles represent significant genes/alleles, orange circles represent genes/alleles of interest from previous screening results).

## METHODS

### *C. elegans* maintenance

*C. elegans* were grown and maintained at room temperature or in incubators at ∼23°C on nematode growth media (NGM) plates seeded with *Escherichia coli* (OP50) as a food source. All strains contained either *him-8(e1489)* or *him-5(e1490)* to increase the number of male animals. Strains, alleles, and transgenes included are listed in **Table 1** ordered by figure number. For most experiments, only male *C. elegans* were analyzed as DVB plasticity does not occur in control hermaphrodites^39^. Hermaphrodites were used for breeding, strain maintenance, and experiments where indicated. In order to stage animals across development and adulthood, which is listed for each experiment, animals were identified as L1 or L4 animals based on size and morphology (body and vulva or male tail tip) and aged to the indicated time of experiment or analysis.

### Confocal Microscopy

10-20 L4 males were picked onto clean NGM plates seeded with OP50 and allowed to age to the indicated day of adulthood. Where indicated for analysis of larval animals, 10-20 L1 animals were picked in the same manner, and for adult hermaphrodite analysis 10-20 L4 hermaphrodites were picked. Animals were anesthetized in 5 μl of 100 mM sodium azide solution on a thin pad of 5% agarose and covered with a glass coverslip. All animals present were analyzed by fluorescence microscopy using an inverted Leica TCS SP8 laser-scanning microscope operated by LAS X software. Identical laser and detector settings were used for all experimental conditions using 63x objective with glycerol immersion fluid (type G). Z-stacks were set such that imaging spanned the entire neuron at a slice size of 0.6μM. Z-stacks were approximately 30-75 slices. Confocal micrographs were generated by compressing z-stacks as maximum intensity projections in FIJI. Representative z-stacks for figures were cropped, analyzed, and arranged in Adobe Photoshop and Illustrator.

### Analysis of DVB neuron morphology

DVB neuron morphology analysis was performed using FIJI as previously described^39–41^, with experimenter blinded to genotype. Analysis was performed using the Simple Neurite Tracer or updated SNT Neuroanatomy plugin. Neurites were traced from the center of the DVB cell soma to the longest point of neurite extension. Branches were added connecting to the existing trace, including all branches posterior to where the axon turns anteriorly into the ventral nerve cord. The Analyze Skeleton functions was used to quantify neurite length and number of junctions (as a proxy for number of branches).

### Analysis of DVB *cla-1::gfp* pre-synaptic puncta

Synaptic *cla-1::gfp* puncta were analyzed using the FIJI Analyze Particle function. Only CLA-1::GFP puncta overlapping with DVB neurites (labeled with wCherry (worm optimized mCherry)) were quantified, using the same neurites of DVB used for neurite tracing. The number of puncta and total puncta size area were quantified. The number of puncta was verified manually to ensure consistency between the observer and the analysis software. Z-stacks were opened in FIJI, and converted to 8-bit and auto-thresholded. If auto-thresholding was not representative of *cla-1::gfp* puncta, small manual adjustments were made to match original images. A region of interest (ROI) was defined to restrict puncta counts to DVB neurites (exclusion of soma and PVT neuron). Pixel size of *cla-1::gfp* puncta was restrained to 0.01 pixels to exclude noise and circularity was not restricted.

### Morphologic variability in the DVB and RIC neurons

We utilized the Python package CAJAL (v1.0)^52^ to analyze 118 neuron morphology reconstructions, of which 39 were DVB at age 1 day, 36 were RIC, and 43 were DVB at age 3 days. For each neuron we sampled 100 points uniformly throughout the skeleton and computed the geodesic pairwise distance matrix between sampled points. We then computed the Gromov-Wasserstein distance between each pair of neurons and created normalized histograms of the distributions of pairwise distances within the RIC class, DVB day 1 class, DVB age 3 class, and between neurons in the RIC and the DVB classes. We tested for differences between the median of the distributions using a two-sided Wilcoxon rank-sum test.

### CAJAL analysis of DVB morphology in conserved autism-associated gene alleles

We utilized the Python package CAJAL (v1.0)^52^ to analyze 1,910 neuron morphology reconstructions. 1,183 of these represented one of 28 alleles under investigation, the remaining 717 were control DVB neurons. We computed the Gromov-Wasserstein distance between each pair of neurons using the same parameters as in the paragraph “*Morphological variability in the DVB and RIC neurons*”. For each allele, we applied the graph Laplacian permutation test implemented in CAJAL^52^ to the set of neurons carrying the allele and the control group. This allowed us to identify alleles that lead to morphological changes with respect to the control group. The false discovery rate was controlled using the Benjamini-Hochberg procedure.

The Gromov-Wasserstein space was then embedded into the Euclidean plane using the multidimensional scaling (MDS) algorithm as implemented in the scikit-learn Python package. For each significant allele, we plotted a circle with radius 𝜎̂/2 centered at the medoid of the cluster, where we defined the sample variance of the allele in the Gromov-Wasserstein space as

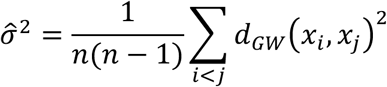

### Aldicarb spicule protraction assay

Spicule protraction behavior was assayed using an aldicarb assay as previously described^39–41^. Aldicarb testing plates were prepared fresh by adding 130 μl of a 100 mM aldicarb in 70% ethanol stock solution to the surface of an NGM plate (7-10 ml) and allowed to dry for one hour. ∼10 males were transferred to the aldicarb plate, and the time was recorded for each animal when spicule protraction of longer than 5 seconds occurred. After spicule protraction, males were removed from the plate. Investigators were blinded to genotype and other experimental conditions.

### Statistics and reproducibility

The number for each experiment was based on previous studies and effect size, with each experiment performed with at least 3 independent replicates and each trial performed with matched controls. Experimenter was blinded to genotypes and conditions for all experiments. All data were analyzed and plotted in GraphPad Prism 9 and statistical significance was determined using one-way ANOVA with Tukey’s post-hoc test. For comparisons of two data sets, a two-tailed unpaired t-test was used to compare significance. “n” represents the number of individual animals tested and is indicated in each figure. Error bars on figures represent standard error of the mean (SEM) and p-values are shown in each figure to indicate significance (P<0.05).

## ACKNOWLEDGEMENTS

The authors thank the labs of Colin C. Conine, David M. Raizen, John I. Murray, and Meera V. Sundaram for their feedback on this project. We also thank Theodore G. Drivas and other members of the Hart lab for support and comments on the manuscript. Some strains were provided by the CGC, which is funded by NIH Office of Research Infrastructure Programs (P40OD010440). This work was supported by a SFARI Bridge to Independence Award (MPH), Penn CEET pilot grant supported by National Institutes of Health grant (P30ES013508)(MPH), National Institutes of Health grant 1R01NS129736 (MPH), National Institutes of Health grant RF1MH130553 (PGC), and NDCN Collaborative Pilot Award from the Chan Zuckerberg Initiative (PGC)

## Author Contributions

KZ and MPH conceived and designed the study and experiments. KZ conducted most behavioral and microscopy experiments and supervised a subset of experiments performed by SV, VT, AFEC, and PT. KZ and MPH generated strains and performed genetic studies. KZ and MPH processed and analyzed data and KZ and MPH interpreted all data. PGC and PN analyzed data with CAJAL. MPH wrote the manuscript and assembled tables and figures with assistance from KZ, PGC, and PN. All authors reviewed, revised, and approved the manuscript.

## Data and materials availability

All data and materials are available upon request to the corresponding author. All data are represented in the main text or the supplemental materials.

