## Supplementary figures and images for "Multiple autism genes influence GABA neuron remodeling via distinct developmental trajectories"

### Supplementary Figures

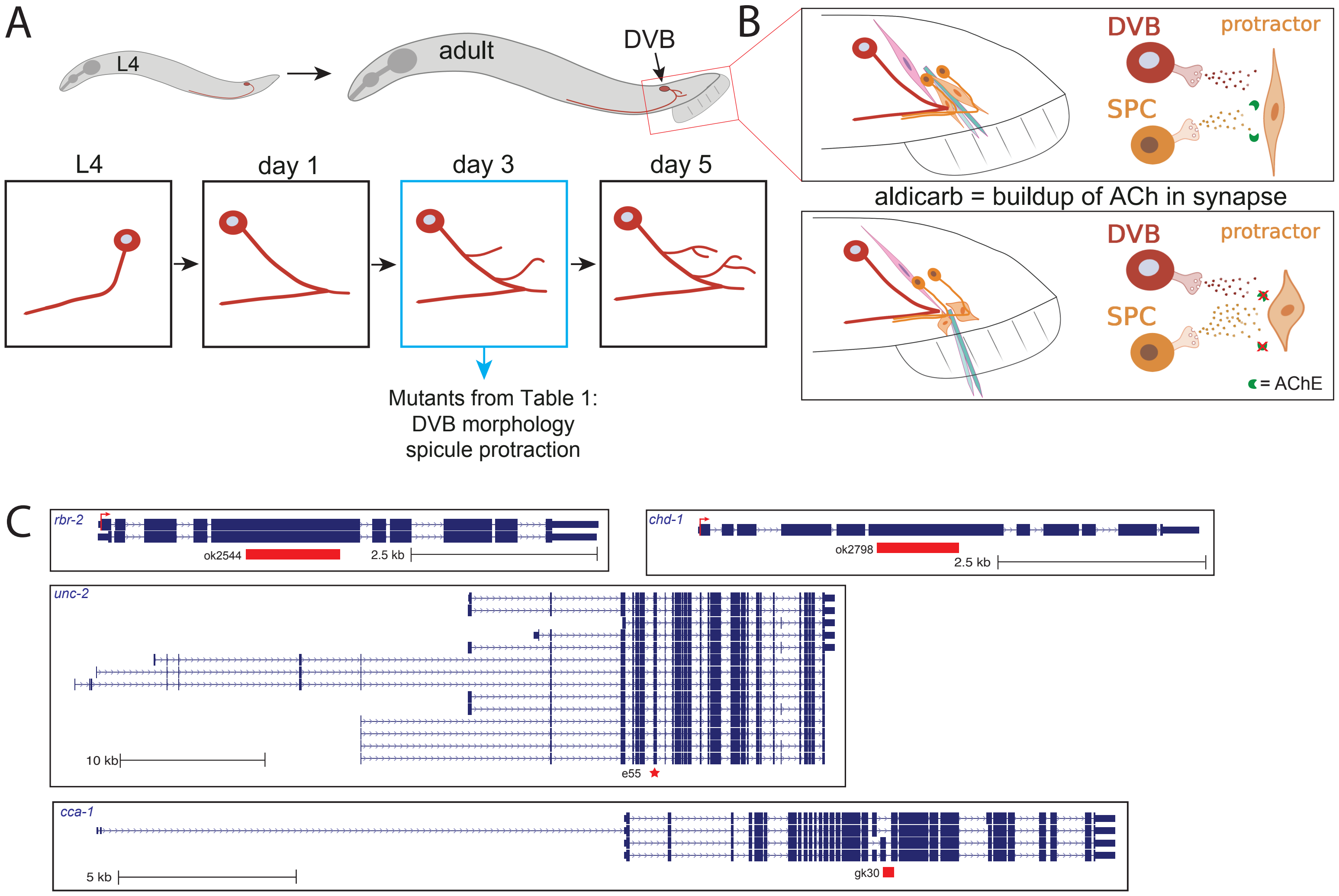


Supplemental Figure 1

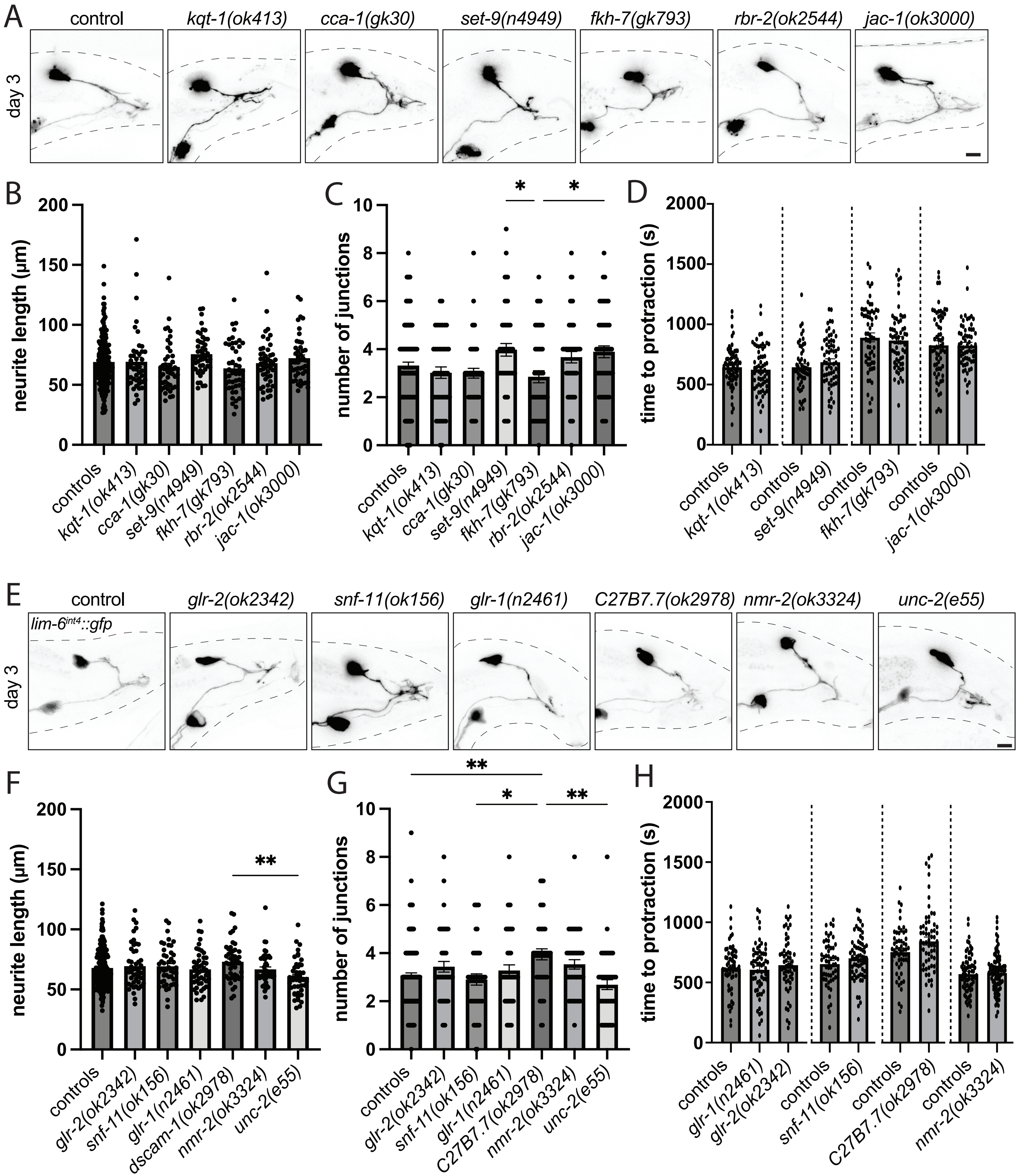


Supplemental Figure 2

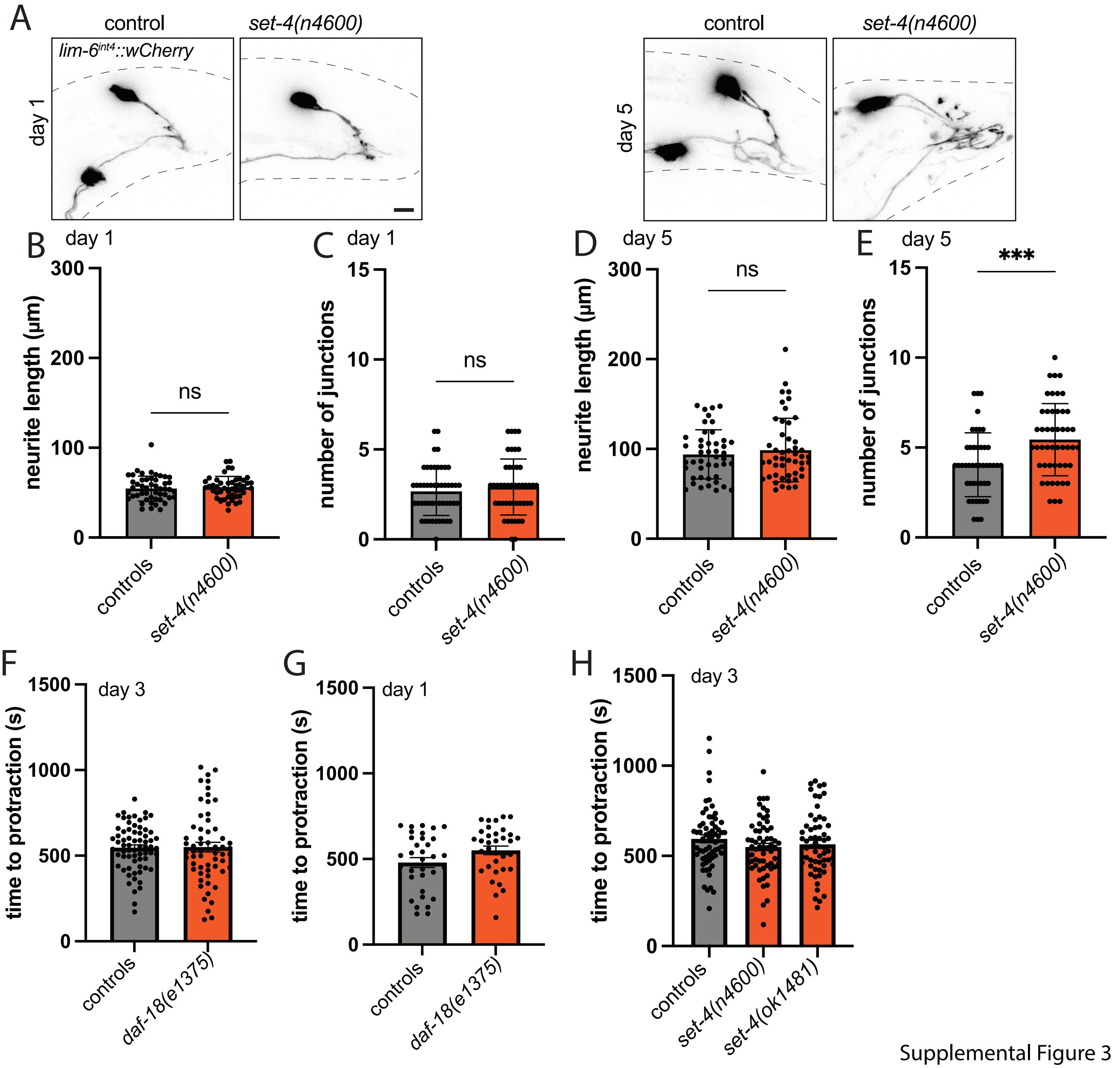


Supplemental Figure 3

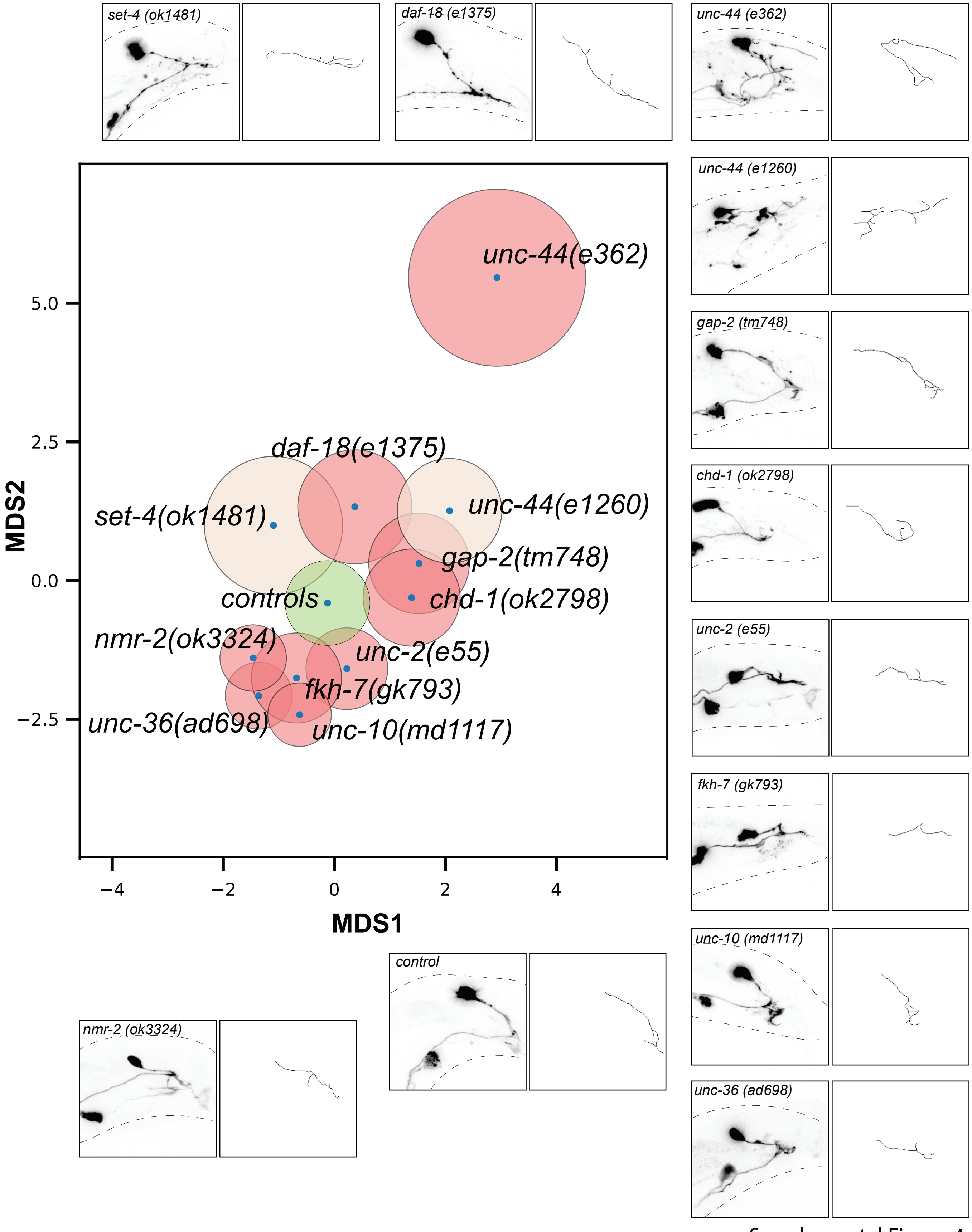


Supplemental Figure 4
